# Identification of *CNTN2* as a genetic modifier of PIGA-CDG through pedigree analysis of a family with incomplete penetrance and functional testing in *Drosophila*

**DOI:** 10.1101/2024.08.12.607501

**Authors:** Holly J. Thorpe, Brent S. Pedersen, Miranda Dietze, Nichole Link, Aaron R. Quinlan, Joshua L. Bonkowsky, Ashley Thomas, Clement Y. Chow

## Abstract

Loss of function mutations in the X-linked *PIGA* gene lead to PIGA-CDG, an ultra-rare congenital disorder of glycosylation (CDG), typically presenting with seizures, hypotonia, and neurodevelopmental delay. We identified two brothers (probands) with PIGA-CDG, presenting with epilepsy and mild developmental delay. Both probands carry *PIGA^S132C^*, an ultra-rare variant predicted to be damaging. Strikingly, the maternal grandfather and a great-uncle also carry *PIGA^S132C^*, but neither presents with symptoms associated with PIGA-CDG. We hypothesized genetic modifiers may contribute to this reduced penetrance. Using whole genome sequencing and pedigree analysis, we identified possible susceptibility variants found in the probands and not in carriers and possible protective variants found in the carriers and not in the probands. Candidate variants included heterozygous, damaging variants in three genes also involved directly in GPI-anchor biosynthesis and a few genes involved in other glycosylation pathways or encoding GPI-anchored proteins. We functionally tested the predicted modifiers using a *Drosophila* eye-based model of PIGA-CDG. We found that loss of *CNTN2*, a predicted protective modifier, rescues loss of *PIGA* in *Drosophila* eye-based model, like what we predict in the family. Further testing found that loss of *CNTN2* also rescues patient-relevant phenotypes, including seizures and climbing defects in *Drosophila* neurological models of PIGA-CDG. By using pedigree information, genome sequencing, and *in vivo* testing, we identified *CNTN2* as a strong candidate modifier that could explain the incomplete penetrance in this family. Identifying and studying rare disease modifier genes in human pedigrees may lead to pathways and targets that may be developed into therapies.

## Introduction

Rare Mendelian genetic disorders are often caused by variants in a single causative gene. However, genetic background plays a large role in the severity and clinical presentation, complicating the genetic landscape^1^. Genetic modifiers are genes that affect a specific disease-causing variant, without causing a disease itself, by either enhancing or suppressing a phenotype^1^. One of the best ways to identify modifier genes in rare disease is to identify pedigrees that show incomplete penetrance or reduced expressivity. Affordable whole genome sequencing allows for rapid nomination of candidate variants underlying phenotypic differences in a pedigree. However, as is often the case, these pedigrees are rare, and in order to go beyond a list of candidates, we need to rapidly test hypotheses in *in vivo* models. This study is one of the first to leverage genetic information from a pedigree showing incomplete penetrance of a rare Mendelian disorder to identify candidate genetic modifiers and test them for genetic interaction *in vivo* in a *Drosophila* model.

Phosphatidylinositol glycan class A congenital disorder of glycosylation (PIGA-CDG) is an ultrarare, X-linked recessive disorder^2,3^. Patients typically present with seizures, hypotonia, and neurodevelopmental delay^2,4^. However, because of the ubiquity of *PIGA* expression, symptoms can occur in any system across the body. There are fewer than 100 reported PIGA-CDG patients, and there is a large amount of phenotypic variability among patients^2^. The pathogenic mechanisms driving symptoms in PIGA-CDG are not understood, and current treatments of PIGA-CDG are limited to symptom-specific treatments. While these may alleviate some of the patients’ symptoms, they do not target the underlying causes of PIGA-CDG.

PIGA-CDG is caused by loss of function mutations in the PIGA gene^2,5^. *PIGA* is an X-linked gene that encodes a protein necessary for synthesizing glycosylphosphatidylinositol (GPI) anchors^4^. GPI anchors are glycolipids used to tether roughly 150 different proteins to the cell surface. GPI anchors are synthesized in the ER and attached to a protein. The resulting glycoprotein is then sent to the Golgi for remodeling before it is sent to the cell membrane. PIGA localizes to the cytoplasmic side of the ER membrane, where it acts as the catalytic component of the N-acetylglucosamine-transferase (GlcNAc-T) complex^6^. GlcNAc-T performs the first step in GPI anchor synthesis by transferring a GlcNAc onto a phosphatidylinositol molecule.

PIGA was initially studied in the context of a somatic mutation in hematopoietic cells, resulting in paroxysmal nocturnal hemoglobinuria^4^. The first case of PIGA-CDG was published in 2012^7^. Initial development of PIGA-CDG models proved difficult because full body knockout of PIGA in mice is embryonic lethal^8^. Cell models from mice and humans are viable and have decreased GPI-anchored proteins on the cell surface^9,10^. However, cell models are limited because many GPI-anchored proteins are involved in cell-cell communication, and those relationships are difficult to reproduce in cell culture^11^. Conditional knockout models in the mouse nervous system are viable and able to recapitulate patient phenotypes such as ataxia and degenerative tremors^10^. Neuron- and glia-specific knockdown of *PIGA* in *Drosophila* also reproduces patient phenotypes, including motor dysfunction and seizures, respectively^12^. Heterozygous knockout of *PIGA* in *Drosophila* also results in severe seizures^12^.

PIGA-CDG patients present with extensive phenotypic variability affecting severity and symptom type^4^. The etiology of the phenotypic variability is poorly understood and may include effects by the specific disease-causing variant, environmental factors, or genetic modifiers. Identifying genetic modifiers of rare diseases like PIGA-CDG is challenging because there are not enough patients to provide statistical power^2^.

We identified an extended, three-generation family showing incomplete penetrance for PIGA-CDG. Using whole genome sequencing from the probands and unaffected carriers in this family, we screened for potential candidate modifier genes. These candidate genes were identified based on inheritance pattern (protective in the unaffected carriers or susceptibility in the probands), effects on glycosylation/GPI-anchor synthesis, and predicted impact of the variants on a protein.

With this pipeline, we identified eight potential protective modifier genes, including a null variant in *CNTN2*, and twelve potential susceptibility genes. Testing for genetic interaction with *PIGA* in a *Drosophila* eye model revealed *CNTN2* as a strong protective genetic modifier, rescuing an eye size defect caused by loss of *PIGA*. *CNTN2* is a GPI-anchored protein involved in neuron-glial cell interaction. Further, we found that loss of *CNTN2* can rescue climbing defects and seizure phenotypes caused by loss of *PIGA* in the neurons and glia, respectively. Loss of *CNTN2*, a proposed protective modifier in the pedigree, rescued multiple phenotypes in *Drosophila* resulting from loss of *PIGA*.

## METHODS

### Consent and clinical data

This study was approved by the University of Utah Institutional Review Board, protocol 40092. Informed consent, assent, or parental permission (as applicable) was obtained from the participants. Clinical data (presence of epilepsy) was obtained through history of the participant by a member of the study team (JLB).

### Whole-genome sequencing

Whole genome sequencing (WGS) was performed after DNA extraction from a venous blood draw. Sequencing was performed at the University of Utah High-Throughput Genomics (HTG) core using the Illumina NovaSeq 6000 platform to sequence 150bp paired-end reads using standard protocols. Whole-genome analysis was performed by the Utah Center for Genomic Discovery (UCGD) at the University of Utah. Sequences were aligned to the GRCh38 human reference genome with Burrows-Wheeler Alignment (BWA-MEM)^13^. Deduplication was performed using SAMBLASTER^14^. Indel realignment, base recalibration, and variant calling were done using Sentieon’s DNAseq variant calling workflow^15^.

### Variant prioritization

We used Slivar^16^ to identify variants following the inheritance modes (Protective and Susceptibility) in CDG genes and GPI-anchored proteins with a GQ > 5. Variants were considered protective if the carriers were heterozygous or homozygous for an alternate allele, while the probands were homozygous for the reference allele (hg38), or if the carriers were homozygous for an alternate allele while the probands were heterozygous. Variants were considered susceptibility variants if the carriers were homozygous for the reference allele while the probands were heterozygous or homozygous for an alternate allele, or if the carriers were heterozygous for an alternate allele while the probands were homozygous for an alternate allele. BED files with genomic coordinates from UniProt were used to identify variants in CDG and GPI-anchored protein genes^17^. Variants categorized as ‘LOW’ and ‘MODIFIER’ impact during the initial variant calling were removed. Missense variants were characterized with SIFT^18^ as either ‘Tolerated’ or ‘Deleterious’ and PolyPhen2^19^ as ‘Benign’, ‘Possibly Damaging’, or ‘Probably Damaging’. Variants that were both ‘Tolerated’ and ‘Benign’ were eliminated.

### Code for variant calling

Slivar code to call protective or susceptibility variants in CDG and GPI-anchored protein genes:

~~~
slivar expr \
     --region cdg_gpi.bed \
     --info “INFO.impactful && variant.FILTER == ‘PASS’” \
     -g gnomad.hg38.genomes.v3.fix.zip \
     --vcf PIGA.vcf.gz \
     -a A631.alias \
     -o susceptibility_cdg_gpi.vcf \
     --pass-only \
     --group-expr “susceptibility:P0CARRIER.hom_ref && P0GPA.hom_ref && AFF1.het && AFF2.het || P0CARRIER.hom_ref && P0GPA.hom_ref && AFF1.hom_alt && AFF2.hom_alt || P0CARRIER.het && P0GPA.het && AFF1.hom_alt && AFF2.hom_alt && P0GPA.GQ > 5 && P0CARRIER.GQ > 5 && AFF1.GQ > 5 && AFF2.GQ > 5”
~~~

~~~
     slivar expr \
     --region cdg_gpi.bed \
     --info “INFO.impactful && variant.FILTER == ‘PASS’” \
     -g gnomad.hg38.genomes.v3.fix.zip \
     --vcf PIGA.vcf.gz \
     -a A631.alias \
     -o protective_cdg_gpi.vcf \
     --pass-only \
     --group-expr “protective:P0CARRIER.het && P0GPA.het && AFF1.hom_ref && AFF2.hom_ref || P0CARRIER.hom_alt && P0GPA.hom_alt && AFF1.hom_ref && AFF2.hom_ref || P0CARRIER.hom_alt && P0GPA.hom_alt && AFF1.het && AFF2.het && P0GPA.GQ > 5 && P0CARRIER.GQ > 5 && AFF1.GQ > 5 && AFF2.GQ > 5”
~~~

cdg_gpi.bed file included as Data S1 supplemental file. Slivar v0.1.12 used for variant calling. VCF and alias files available upon request.

### Fly stocks and maintenance

Stocks were maintained on standard Archon Scientific glucose fly food at 25°C on a 12-h light/dark cycle. The *w-;; eya composite-GAL4* and *w-;eya composite-GAL4/CyO;+* lines were a gift from Justin Kumar (Indiana University Bloomington) and were characterized previously^20^. Repo-GAL4 (7415) and elav-GAL4 (46655) were from Bloomington *Drosophila* Stock Center. *Drosophila* orthologues of candidate genes were determined using DIOPT, requiring a score ≥6^21^. RNAi lines used in this study were ordered from the Bloomington Drosophila Stock Center and the Vienna Drosophila Resource Center. The PIG-A null allele was generated as part of the Gene Disruption Project. The PIGA-null allele was made as a PIG-A Kozak-fullGAL4 allele (CR70080)^22^ using the sgRNAs TATGTGGTATGTCAAATTACTGG and TTGGCTAGTTGATGGAAGAATGG.

To generate the *dPIGA^WT^* and *dPIGA^S100C^* transgenes, *PIGA* coding sequence from BDGP DGC clone LD44262 was transferred to pGW-HA.attB using NEB Gibson Assembly master mix (E2611). pGW-HA.attB ccdB region was removed using AgeI and KpnI. *PIGA* was amplified with primers 5’-TCAAAAGGTATAGGAACTTCAACCGGTCAACATGCGCATCTGCATGGTGTCGGAC-3’ and 5’-TATACTAAGTAGAGAATAGGAACTTCGAGGTACctaGCTGGCCCTACGTTGTAC-3’, which include homology regions to pGW-HA.attB. To generate the fly mutation corresponding to S100C, site directed mutagenesis was performed using Q5 Site-Directed Mutagenesis Kit (New England Biolabs E0554) with primers 5’ CTCTGCCTTCtGTGCTCTGGC-3’ and 5’-TGGCCATGGACCACTTCC-3’. Resulting clones were sequenced using long read sequencing technology (Plasmidsaurus), and transgenic animals were generated using ϕC31 mediated site-specific integration into PBac{y[+]-attP-3B}VK00033 (Bestgene).

To test d*PIGA^WT^* or *dPIGA^S100C^* transgene rescue, *w-; PIGA null Kozak-fullGAL4/CyO;UAS-dPIGA^transgene^* flies were created through genetic crosses and self-crossed. Resulting progeny was scored comparing numbers of homozygous *PIGA null* flies to the balancer class heterozygotes. Proportion of expected survival was calculated based on the number of balancer class heterozygotes.

To create double knockdown flies, a model of *w-; PIGA RNAi/CyO;GAL4/TM3* was crossed to either the candidate modifier RNAi lines or the equivalent control lines (*attP40, attP2, attP, w1118)* selecting progeny that were negative for both *CyO* and *TM3, Sb* balancers.

### Phenotypic analysis

#### Eye imaging and quantification

Crosses were performed initially performed at 25°C. Lethality is high among PIGA knockdown flies, so if sufficient flies were unable to be collected, the cross was repeated at 20°C. Adult female flies aged 3-5 days were collected under CO_2_ anesthesia then frozen at -80°C for later imaging. Eyes were imaged at 3x magnification using the Leica EC3 Camera. Eye area was measured as previously described^23^.

#### Negative Geotaxis Assay

Crosses were performed at 25°C. Flies were maintained off CO_2_ for at least one day after collection. Tested flies were 3-5 days of age. Ten flies were placed in empty vials and tapped to drop flies to the bottom. The number of flies able to climb off the bottom of the vial within 10 seconds was counted.

#### Bang Sensitivity Assay

Crosses were performed at 25°C. Flies were maintained off CO_2_ for at least one day after collection. Tested flies were 3-5 days of age. Ten flies were placed in empty vials. Vials were immediately vortexed for 10 seconds, and seizure behavior was observed.

## Statistics

Transgene rescue significance was determined using Student’s T-test in R. *PIGA^WT^* survival significance was calculated using a one-sample t-test in R with mu = 1. *PIGA^S132C^* survival significance was calculated using a two-sample t-test in R comparing *PIGA^S132C^* survival to *PIGA^WT^*.

Genetic interactions between PIGA and proposed modifier genes in the *Drosophila* eye model were determined using a two-way ANOVA run in RStudio. PIGA anchor gene level (wild type or knockdown) and candidate modifier gene level (wild type or knockdown) were the variables compared in the analysis.

Chi-square was performed in Excel using *PIGA* single knockdown proportions to determine expected counts compared to the double knockdown counts.

## Results

### Incomplete penetrance in a family with PIGA-CDG

We identified a family with two brothers diagnosed with PIGA-CDG, presenting with early-onset epilepsy and mild developmental delay (Figure 1A). These probands were previously reported in Bayat et al., 2020 (patients 24 and 25). An epilepsy multi-gene panel identified a hemizygous missense mutation*, PIGA^S132C^* (*PIGA c.395C>G*), in both probands. Diagnosis was confirmed by measuring cell surface levels of GPI-anchored proteins (Figure S1)^2,24^. Probands have ∼50% reduction in cell surface GPI-anchored proteins compared to control levels. The two brothers were consequently diagnosed with PIGA-CDG, although their symptoms are milder than most reported PIGA-CDG patients.

**Figure 1:**
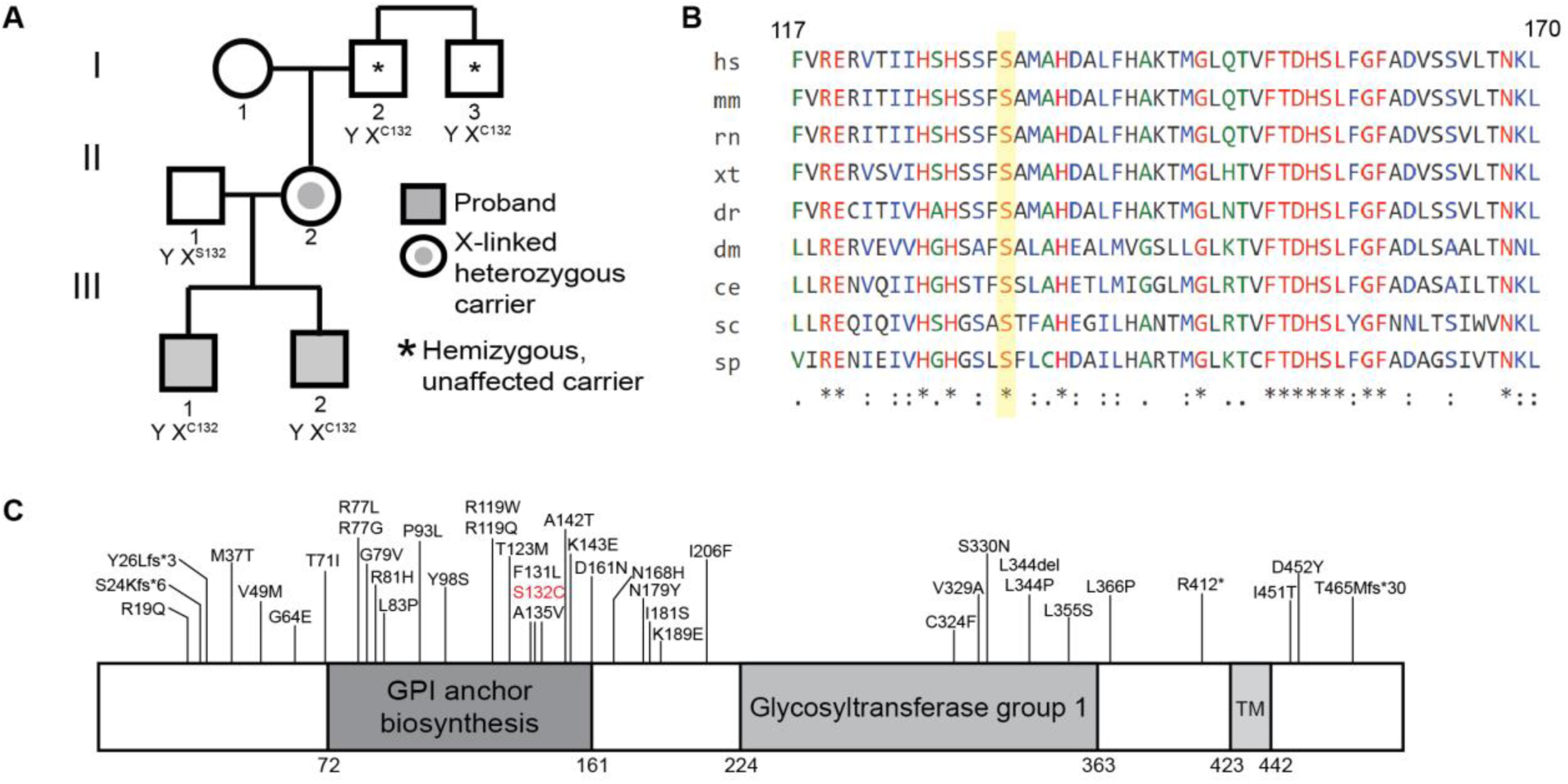
A) PIGA-CDG pedigree with incomplete penetrance. B) S132 (highlighted in yellow) and surrounding amino acids are highly conserved from yeast to humans. (*) indicates completely conserved positions; (:) indicates position is semi-conserved with substitutions showing strongly similar properties; (.) indicates semi-conserved positions with substitution showing weakly similar properties. (sp - *Schizosaccharomyces pombe*, sc - *Saccharomyces cerevisiae*, ce - *Caenorhabditis elegans*, dm - *Drosophila melanogaster*, dr - *Danio rerio*, xt - *Xenopus tropicalis*, rn - *Rattus norvegicus*, mm - *Mus musculus*, hs - *Homo sapiens*). C) Schematic of PIGA protein with domains and known PIGA-CDG causing variants^2^. S132C in red is variant found in probands.

The *PIGA^S132C^* variant is ultra-rare and not found in gnomAD^25^. These probands are the first patients reported with the *PIGA^S132C^* variant. S132 is evolutionarily invariant from yeast to mammals (Cons 100 Verts = 9.564) (Figure 1B)^26^. *PIGA^S132C^* is in the GPI anchor biosynthesis domain of PIGA (Figure 1C). Other disease-causing variants are found in this domain, including at positions 131 and 135. The *PIGA^S132C^* variant was predicted to be Deleterious by SIFT, Probably Damaging by PolyPhen-2, and Disease Causing by MutationTaster^2^.

Further genotyping of the family revealed that the probands inherited the disease variant from their mother (Figure 1C). Since the disorder is X-linked recessive, she is healthy. Strikingly, the healthy mother inherited the variant from her father, the probands’ grandfather. The grandfather’s brother (great-uncle to probands) also carries the *PIGA^S132C^* variant. Both the grandfather and great-uncle are asymptomatic (unaffected carriers). Thus, the *PIGA^S132C^* variant shows incomplete penetrance in this pedigree, suggesting a role of genetic modifiers.

### PIGA^S132C^ is a partial loss-of-function allele

We determined the functional consequences of the *PIGA^S132C^* variant by testing whether transgenic expression can rescue a *PIGA* null allele in a *Drosophila* PIGA-CDG model. Homozygous null *PIGA Drosophila* are unable to survive into adulthood^12^. Using a null *PIGA* allele where the coding sequence is replaced by a GAL4 cassette, we drove expression of two *UAS-PIGA* transgenes. The transgenes either encoded wild-type *Drosophila PIGA* (d*PIGA^WT^*) or *Drosophila PIGA* with the patient variant (*PIGA^S100C^* in *Drosophila*; *dPIGA^S100C^*). We expected the *dPIGA^WT^* transgene to provide complete rescue and the *dPIGA^S100C^* transgene to provide partial rescue, consistent with a partial loss-of-function mutation similar to those found in other PIGA-CDG patients^2,5^. Across three replicates, *dPIGA^WT^* expression provided an average of 86% expected rescue, which was not statistically different from full 100% rescue (p = 0.154) (Figure 2). On average, *dPIGA^S100C^* provided 59% expected rescue, significantly lower than *dPIGA^WT^* (p = 0.039) (Figure 2), indicating that the *PIGA^S132C^* variant is likely a partial loss-of-function allele.

**Figure 2:**
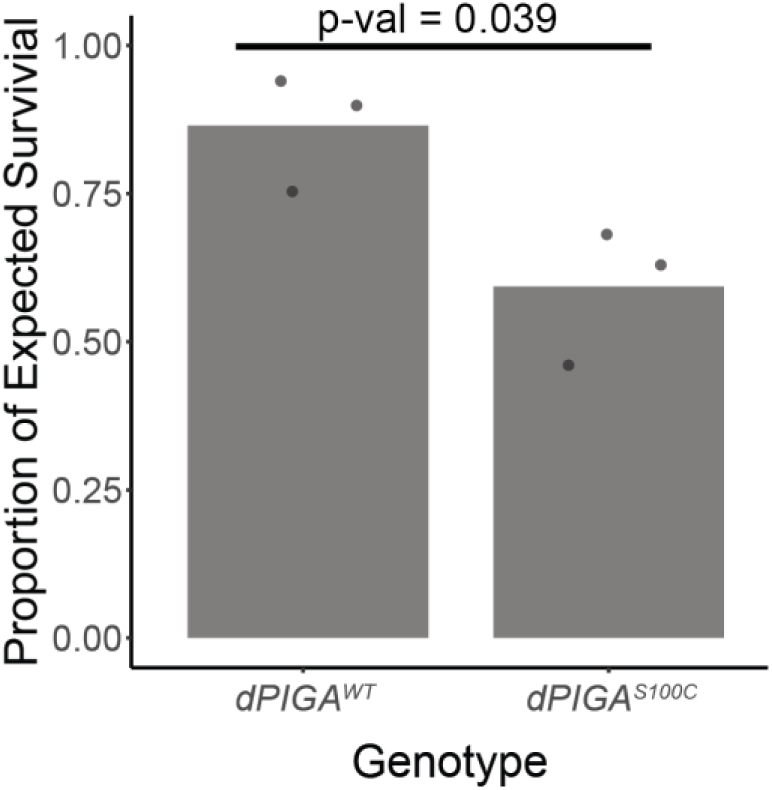
Proportion of PIGA null Drosophila rescued by *dPIGA^WT^* and *dPIGA^S100C^* transgenes. *dPIGA^S100C^* transgene provides partial rescue compared to *dPIGA^WT^* (p = 0.039) indicating variant is partial loss-of-function. *dPIGA^WT^* rescue levels were not significantly different from expected complete 100% rescue (p = 0.154). *dPIGA^WT^* n = 416, 432, 391; *dPIGA^S100C^* n = 174, 154, 134.

### Genome sequencing of family members reveals potential genetic modifiers

There are two possible inheritance modes for genetic modifiers that are likely explain the incomplete penetrance in the family. There could be a susceptibility variant found in both probands, but not in either of the unaffected carriers. Alternatively, there could be a protective variant found in both unaffected carriers that was not passed down to either proband. To identify these candidate variants, we performed whole genome sequence on these four key members of the pedigree. We used Slivar^16^ to identify candidate variants fitting either of these inheritance patterns (Methods). This genome-wide analysis resulted in 751,347 susceptibility variants and 665,135 protective variants (Figure 3, Table S1).

**Figure 3:**
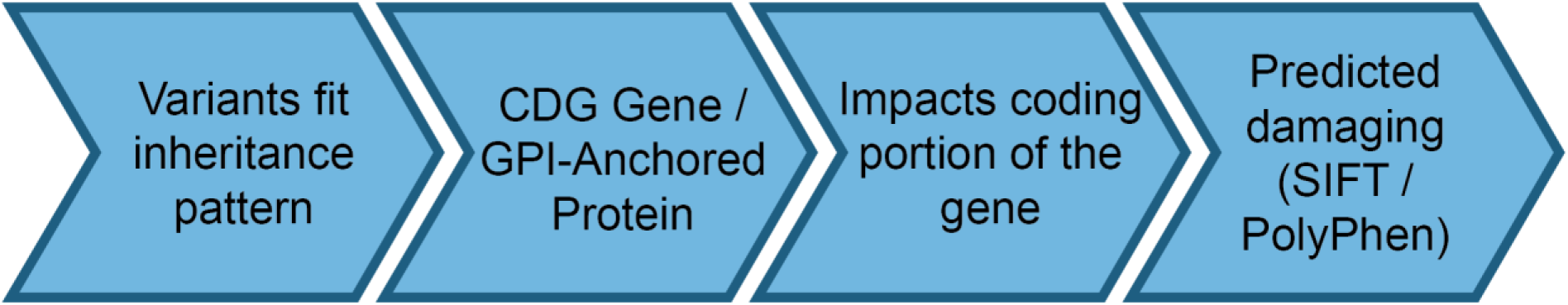
Pipeline to identify most likely candidate variants/genes modifying PIGA-CDG. Number of genes and variants remaining after each criterion found in Table S1.

Previous studies from our lab and others have found that CDG genes are often modifiers of other CDG genes in model organisms and humans^27–31^. Because of this well-known phenomenon, we initially focused our analyses on 189 known CDG genes along with four other genes involved directly with GPI anchor synthesis that have not yet been associated with a CDG (Data S1)^4,32^. We also included 143 known GPI-anchored proteins identified using Uniprot (Data S1)^17^. Using Slivar^16^, we analyzed the protective and susceptibility variants to identify variants in CDG or GPI-anchored protein genes. There were 30 protective variants found in 25 CDG and GPI-anchored protein genes and 30 susceptibility variants found in 27 CDG and GPI-anchored protein genes (Figure 3, Table S1). To further refine the search, we only included variants that impact the coding portion of the genome and were predicted to be either ‘Deleterious’ by SIFT^18^ or ‘Possibly Damaging’ or ‘Probably Damaging’ by PolyPhen-2^19^. This resulted in seven possible protective variants and thirteen possible susceptibility variants (Figure 3, Table S1). All variants are heterozygous, except the variant found in *SPRN* which is heterozygous in the carriers and homozygous in the probands. None of the genes are shared between the two lists.

Among the candidate variants, two protective and one susceptibility variants were in three genes involved in GPI anchor synthesis. Two protective and four susceptibility variants were in six CDG genes not involved in GPI anchor synthesis. Four protective and seven susceptibility variants were in eleven genes encoding GPI-anchored proteins. The three variants affecting GPI anchor synthesis were in *PIGS*, *DPM1*, and *PGAP5* (Table 1). *PIGS* is involved in the GPI transamidase complex that attaches the protein to the synthesized GPI anchor^6^. *DPM1* is involved in the complex that transports mannose into the lumen of the ER. Each GPI anchor requires three to four mannoses. *PGAP5* performs fatty acid remodeling on the GPI anchor, which signals for transport from the ER to the Golgi^33^. The seven variants found in other CDG genes span multiple glycosylation pathways (Table 1). *POMT1*, *LFNG*, and *CRPPA* are involved in O-linked glycosylation^34–36^. *ALG6* is a glucosyltransferase for N-linked glycosylation^37^. *SLC39A8* is a cell surface manganese transporter^38^. Manganese is a necessary cofactor for several galactosyltransferases important in glycosylation. *TRAPPC11* is important for transporting glycoproteins from the ER to the Golgi^39^. The eleven genes encoding GPI-anchored proteins were: *CNTN2, BST1, SPRN, RNT4RL1, SMPDL3B, CD48, ART3, ACHE, MMP17, MDGA2,* and *RGMA* (Table 1).

**Table 1:**
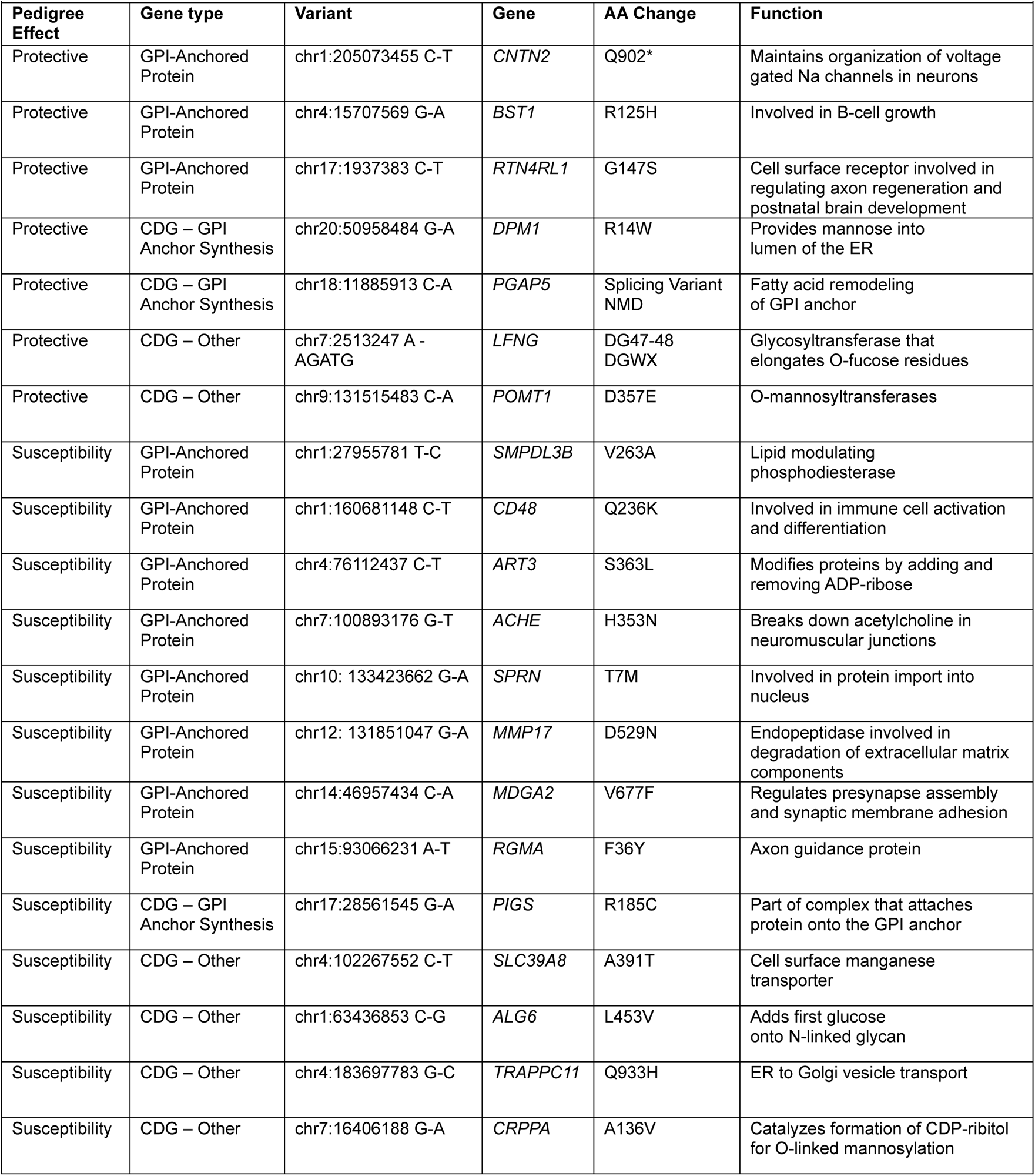
Top candidate modifier variants.

| **Pedigree Effect** | **Gene type** | **Variant** | **Gene** | **AA Change** | **Function** |
| --- | --- | --- | --- | --- | --- |
| Protective | GPI-Anchored Protein | chr1:205073455 C-T | *CNTN2* | Q902* | Maintains organization of voltage gated Na channels in neurons |
| Protective | GPI-Anchored Protein | chr4:15707569 G-A | *BST1* | R125H | Involved in B-cell growth |
| Protective | GPI-Anchored Protein | chr17:1937383 C-T | *RTN4RL1* | G147S | Cell surface receptor involved in regulating axon regeneration and postnatal brain development |
| Protective | CDG – GPI Anchor Synthesis | chr20:50958484 G-A | *DPM1* | R14W | Provides mannose into lumen of the ER |
| Protective | CDG – GPI Anchor Synthesis | chr18:11885913 C-A | *PGAP5* | Splicing Variant NMD | Fatty acid remodeling of GPI anchor |
| Protective | CDG – Other | chr7:2513247 A - AGATG | *LFNG* | DG47-48 DGWX | Glycosyltransferase that elongates O-fucose residues |
| Protective | CDG – Other | chr9:131515483 C-A | *POMT1* | D357E | O-mannosyltransferases |
| Susceptibility | GPI-Anchored Protein | chr1:27955781 T-C | *SMPDL3B* | V263A | Lipid modulating phosphodiesterase |
| Susceptibility | GPI-Anchored Protein | chr1:160681148 C-T | *CD48* | Q236K | Involved in immune cell activation and differentiation |
| Susceptibility | GPI-Anchored Protein | chr4:76112437 C-T | *ART3* | S363L | Modifies proteins by adding and removing ADP-ribose |
| Susceptibility | GPI-Anchored Protein | chr7:100893176 G-T | *ACHE* | H353N | Breaks down acetylcholine in neuromuscular junctions |
| Susceptibility | GPI-Anchored Protein | chr10: 133423662 G-A | *SPRN* | T7M | Involved in protein import into nucleus |
| Susceptibility | GPI-Anchored Protein | chr12: 131851047 G-A | *MMP17* | D529N | Endopeptidase involved in degradation of extracellular matrix components |
| Susceptibility | GPI-Anchored Protein | chr14:46957434 C-A | *MDGA2* | V677F | Regulates presynapse assembly and synaptic membrane adhesion |
| Susceptibility | GPI-Anchored Protein | chr15:93066231 A-T | *RGMA* | F36Y | Axon guidance protein |
| Susceptibility | CDG – GPI Anchor Synthesis | chr17:28561545 G-A | *PIGS* | R185C | Part of complex that attaches protein onto the GPI anchor |
| Susceptibility | CDG – Other | chr4:102267552 C-T | *SLC39A8* | A391T | Cell surface manganese transporter |
| Susceptibility | CDG – Other | chr1:63436853 C-G | *ALG6* | L453V | Adds first glucose onto N-linked glycan |
| Susceptibility | CDG – Other | chr4:183697783 G-C | *TRAPPC11* | Q933H | ER to Golgi vesicle transport |
| Susceptibility | CDG – Other | chr7:16406188 G-A | *CRPPA* | A136V | Catalyzes formation of CDP-ribitol for O-linked mannosylation |

### Candidate modifier genes modify loss of *PIGA in vivo*

Glycosylation is a highly conserved process from yeast to humans^6,40,41^. This conservation makes it possible to test for genetic modifiers of *PIGA* in *Drosophila.* Ubiquitous loss of *PIGA* is lethal in*Drosophila,* making it challenging to test for modifier effects on lethality in a ubiquitous context^12^. Loss of *PIGA* in the *Drosophila* eye results in both quantitative and qualitative defects^27,29,42^. We commonly use the eye as a model to test for disease-relevant genetic interactions^23,27,29,43,44^. Using the GAL4-UAS system, we created a tissue-specific knockdown of *PIGA* with the *eyes absent* GAL4 driver (*eya*) and a *UAS-PIGA* RNAi transgene^20,27^. This combination results in expression of *PIGA* RNAi in the *Drosophila* eye. *PIGA* knockdown in the eye causes a drastic decrease in eye size and leads to glassy eyes and necrosis (Dalton et al., 2022; Thorpe et al., 2024) (Figure 4). We can test for genetic interactions by creating double knockdowns of *PIGA* and the candidate modifiers. Eye size serves as a quantitative readout to determine if the modifier can rescue (make the eye bigger) or enhance (make the eye smaller) the *PIGA* phenotype^23,27,29,43,44^. We tested twelve candidate modifiers for effects on the eye (Table 2, Figure S2, Data S2). *CRPPA*, *CD48, ART3, MMP17, MDGA2, RGMA, BST1, SPRN,* and *RTN4RL1* were excluded because they have no *Drosophila* orthologues.

**Figure 4:**
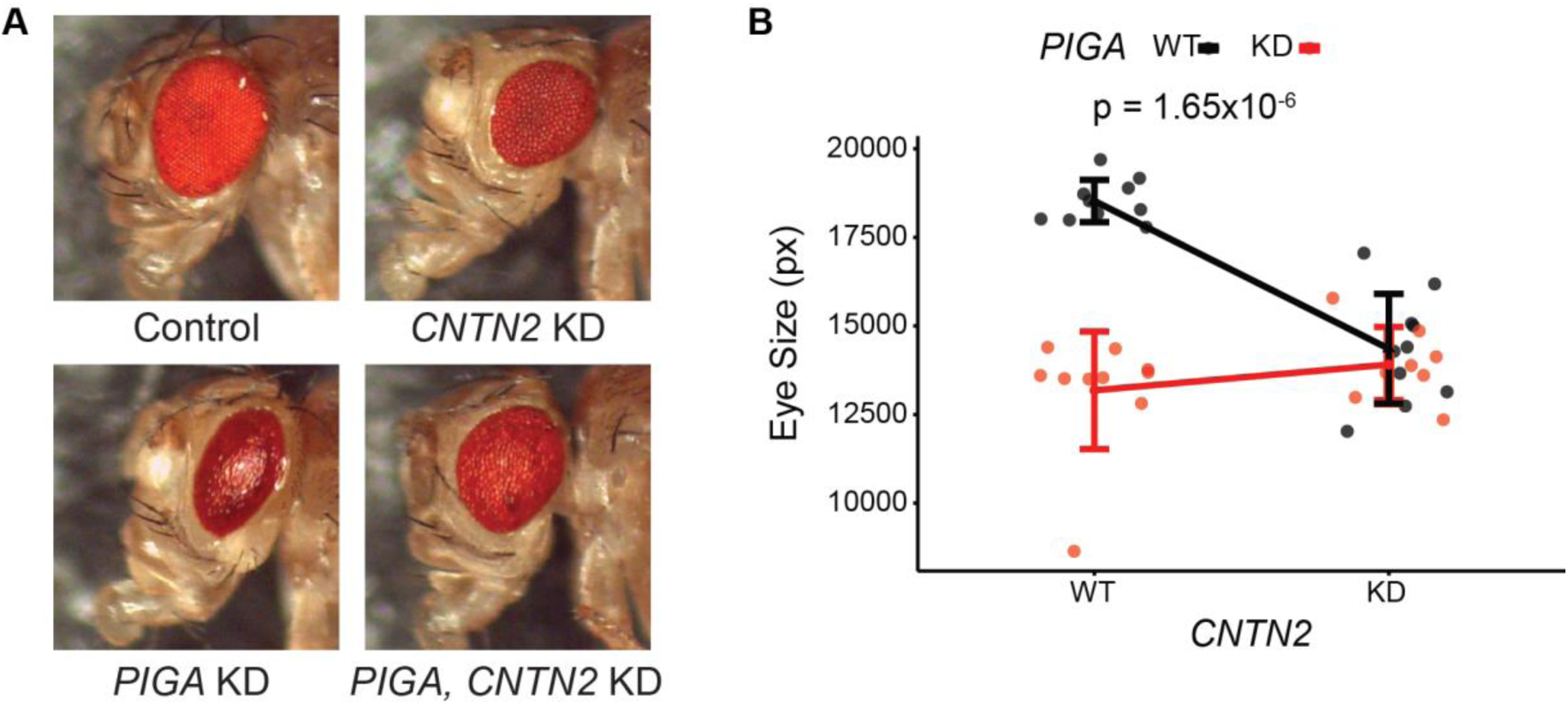
*In vivo* testing for genetic interaction with *PIGA* in the eye. A) Representative images of *Drosophila* eyes showing WT control, single KDs, and double KD eyes. B) Quantification of eye sizes. Quantification in pixels (px) is along the Y axis. X axis is the *CNTN2* genotype: Wild Type (WT) or Knockdown (KD), and line color indicates *PIGA* genotype. Parallel slopes indicate no interaction. Interaction p-value calculated using a two-way ANOVA.

**Table 2:** Effects of knockdown of top candidate modifiers in Drosophila PIGA eye model.

|  |  |  |
| --- | --- | --- |
| **Pedigree effect** | **Gene** | **Effect on *PIGA* eye model** |
| Protective | *CNTN2* | Suppressor |
| Protective | *DPM1* | Enhancer |
| Protective | *PGAP5* | Enhancer |
| Protective | *POMT1* | Enhancer |
| Protective | *LFNG* | Enhancer |
| Susceptibility | *SMPDL3B* | Enhancer |
| Susceptibility | *ACHE* | Suppressor |
| Susceptibility | *PIGS* | Enhancer |
| Susceptibility | *SLC39A8* | Enhancer |
| Susceptibility | *ALG6* | Enhancer |
| Susceptibility | *TRAPPC11* | Enhancer |

Predicted protective modifiers are expected to suppress the *PIGA* phenotype. Of those tested, only *CNTN2* suppressed the eye phenotype caused by loss of *PIGA* (Figure 4, Figure S3). The other four predicted protective modifiers showed significant interaction with *PIGA* (Table 2, Figure S2), but enhanced the eye phenotype caused by loss of *PIGA*, acting in the opposite direction predicted by the pedigree. Predicted susceptibility candidates are expected to enhance the PIGA phenotype. Six of the seven susceptibility candidates tested enhanced the *PIGA* eye phenotype, and one suppressed the phenotype (Table 2, Figure S2). It is not surprising that so many genes enhanced the phenotype, as it is often easier to make a phenotype worse.

We highlight *CNTN2,* encoding a GPI-anchored protein, here and focused our further in-depth analyses on *CNTN2* because the protective variant is a nonsense variant (Q902*), and the eye-based validation data strongly matched the predicted protective effect. The *Drosophila* orthologue for *CNTN2* is *Cont* (DIOPT = 13/16), but herein will be referred to as *CNTN2*. *CNTN2* knockdown in the *Drosophila* eye results in a much milder eye phenotype, with a smaller decrease in eye size compared to *PIGA* and results in minor disorganization of the ommatidia (Figure 4, Figure S3). Surprisingly, double knockdown of *PIGA* and *CNTN2* results in an eye that resembles the much less severe *CNTN2* single knockdown eye in both size and quality, demonstrating a genetic interaction between the two genes (p = 1.65x10^-6^) (Figure 4). In other words, loss of *CNTN2* partially rescues loss of *PIGA*. If the action of both genes was additive, we would observe an eye phenotype that is even more severe than *PIGA* alone. A similar rescue was observed using a second RNAi line for *CNTN2* (p = 2.35x10^-8^) (Figure S4).

### Loss of *CNTN2* rescues *PIGA* neurological phenotypes

Many PIGA-CDG patients experience muscular disorders such as hypotonia and muscle weakness^45^. We previously demonstrated that these symptoms can be recapitulated in neuron-specific *PIGA* knockdown in *Drosophila*^12^. Neuron-specific *PIGA* knockdown results in a movement disorder, as measured by climbing in the negative geotaxis assay^12^. We used the pan-neuronal driver *Elav-gal4* to drive UAS-RNAi against *PIGA* and *CNTN2* to make neuron-specific knockdowns. Then we performed a negative geotaxis assay to determine whether loss of *CNTN2* can also rescue *PIGA* climbing defects.

*PIGA* neuronal knockdown flies are unable to climb to the top of a vial in the same amount of time as wild-type flies^12^. In this study, only 4% of *PIGA* knockdown flies are able to climb off the bottom of a vial in 10 seconds (Figure 5A, Video S1). *PIGA* knockdown flies never make it to the top. *CNTN2* neuronal knockdown resulted in 76% of flies being able to climb off the bottom of the vial in 10 seconds (Figure 5A, Video S2). Similar to what we observed in the eye, double knockdown flies showed improved climbing ability as compared to single PIGA knockdown, with 18% climbing off the bottom of the vial (Figure 5A, Video S3). Knockdown of *CNTN2* partially rescues the climbing ability of single *PIGA* knockdown, demonstrating a genetic interaction between the two genes (p = 2.07x10^-3^).

**Figure 5:**
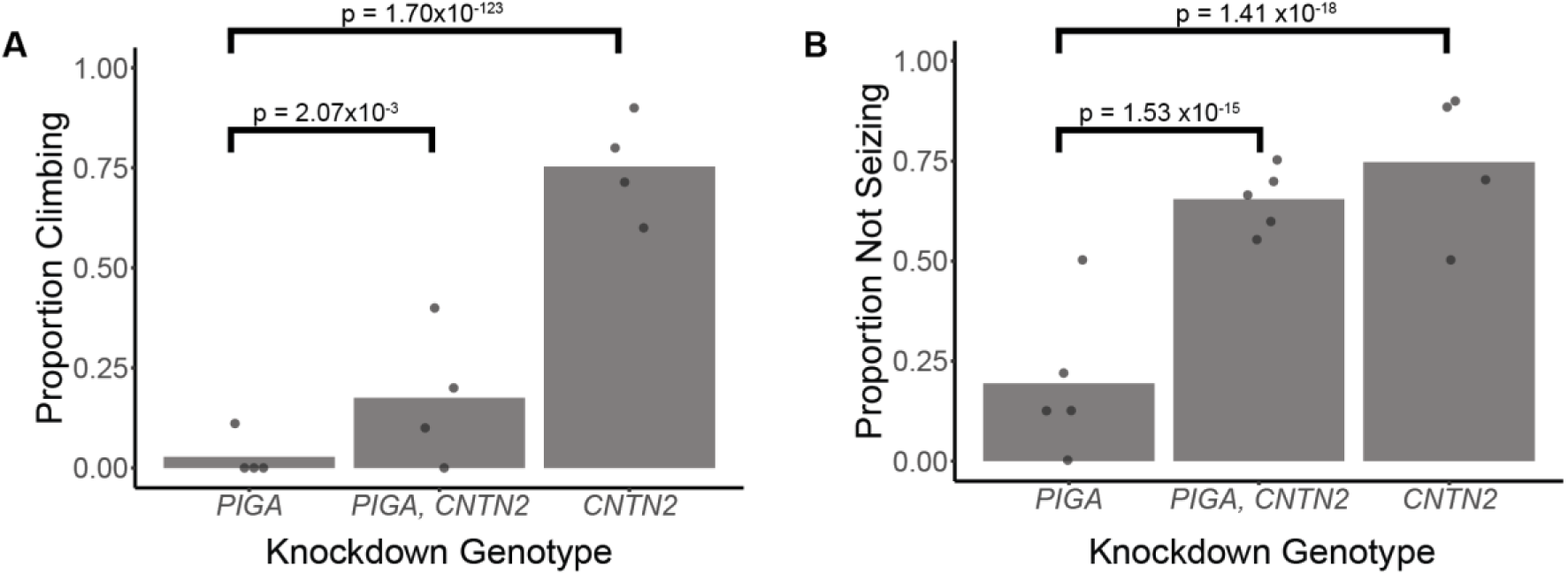
*In vivo* testing for genetic interaction with between *PIGA* and *CNTN2* in neurological models. A) Proportion of flies climbing in single and double neuronal-specific knockdown flies. *PIGA* KD n = 5, 6, 8, 9; *CNTN2* KD n = 7, 10, 10, 10; *PIGA, CNTN2* KD n = 10, 10, 10, 10. B) Quantification of non-seizure like behavior in single and double glial-specific knockdown flies. *PIGA* KD n = 8, 8, 8, 9, 9; *CNTN2* KD n = 9, 10, 10, 10; *PIGA, CNTN2* KD n = 8, 9, 9, 10, 10. P-values calculated using a Chi-square test.

Previous studies from our lab also established that knockdown of *PIGA* in glial cells results in seizures in *Drosophila*^12^. Because seizures are prevalent in PIGA-CDG patients, including the probands in this study, we used this model to test the ability of *CNTN2* to modify seizure phenotypes. We used the pan-glial driver, *repo-GAL4*, to drive UAS-RNAi against *PIGA* and *CNTN2* to create glial-specific knockdowns. We performed a bang-sensitivity assay to measure seizures in the different models. Typical *Drosophila* seizures are characterized by seizure-like spasms, followed by paralysis, then a recovery seizure, and finally, a refractory period when they are resistant to further seizures before they completely recover^46^.

Glial-specific knockdown of *PIGA* results in 81% of flies showing seizure-like activity (Figure 5B, Video S4). During these seizures, the majority of flies experience complete paralysis, and many take over 5 minutes to recover, representing a very severe seizure phenotype. Knockdown of *CNTN2* in glia results in 26% of flies showing very mild seizure-like activity (Figure 5B, Video S5). These flies experience sporadic, fluttering movement, but no paralysis. Based on the effects of each of the single knockdowns, if there is no genetic interaction between *PIGA* and *CNTN2*, the double knockdown flies should show very severe seizures, with nearly 100% of flies displaying seizure-like activity and few, if any, flies showing normal behavior. Again, as we observed with the eye and climbing defects, the double knockdown was much less severe than the single *PIGA* knockdown, with 65% of the double knockdown flies showing normal non-seizure-like behavior and only 35% of flies showing seizure-like activity (Figure 5B, Video S6). *PIGA* and *CNTN2* show a robust genetic interaction (p = 1.53x10^-15^). The seizures in the double knockdown are qualitatively similar to the *CNTN2* single knockdown, with most flies having mild sporadic movements and only occasional paralysis. Loss of *CNTN2* rescues the loss of *PIGA* in glia and reduces the amount and severity of seizures.

## Discussion

The pathophysiology of PIGA-CDG is poorly understood and consequently lacks treatment options^47^. Identifying genetic modifiers of *PIGA* can help to elucidate the pathology of PIGA-CDG and identify therapeutic targets. Here, we used whole genome sequencing from a family showing incomplete penetrance of PIGA-CDG to identify potential genetic modifiers of *PIGA*. *In vivo* testing in *Drosophila PIGA* models showed genetic interactions between *PIGA* and multiple candidate modifiers. *CNTN2* stood out as a protective modifier that was able to rescue phenotypes from loss of *PIGA* in eye tissue, neurons, and glia.

*CNTN2* knockdown rescued loss of *PIGA* in multiple *Drosophila* models of PIGA-CDG. *CNTN2* encodes a GPI-anchored protein^48^. *CNTN2* is predominantly expressed in neuronal tissues^49^. CNTN2 is important for neuronal-glial cell-cell interactions^50^. It works in a network of proteins to attach the myelin sheath to the axon. CNTN2 also aids in the organization of ion channels along the axon. Interestingly, mutations in *CNTN2* have also been associated with an autosomal recessive inherited epilepsy^51,52^. In PIGA-CDG, the partial loss of function of *PIGA* results in decreased synthesis of GPI anchors^2^. It is possible that reducing levels of *CNTN2* alleviates demand on an already stressed GPI anchor synthesis pathway in PIGA-CDG. This general mechanism has been observed in other studies of CDG modifiers where reducing flow through other glycosylation pathways can rescue phenotypes resulting from loss of function in a CDG gene^27,30^. It is also possible that *CNTN2* modifies *PIGA* through its function in neuronal signaling. Further testing is needed to determine the mechanism of interaction.

It remains uncertain whether *CNTN2* is a specific modifier for the *PIGA^S132C^* variant or a general modifier of PIGA loss-of-function. However, the fact that we show improvement in phenotypes across different tissue contexts, using RNAi against *PIGA,* suggests that this is a general mechanism and not specific to the S132C allele. In the future, it will be useful to generate a series of allele-specific models in *Drosophila* to test whether *CNTN2* can be applied to a variety of pathogenic *PIGA* variants.

Our screen also identified *ACHE* as a potential modifier of PIGA-CDG. In the pedigree, it was predicted to be a susceptibility modifier, but it provided a partial rescue in the PIGA eye model. ACHE is also a GPI-anchored protein^53^. In *Drosophila,* it is always GPI-anchored; however, in humans, it is only GPI-anchored in the blood^54,55^. The rescue in the eye model caused by both *CNTN2* and *ACHE* could indicate that reducing GPI-anchored proteins could reduce demand on the GPI anchor synthesis pathway and be a general mechanism of rescue in GPI anchor synthesis disorders. More work is needed to determine if reducing GPI-anchored protein levels may be a promising therapeutic approach for PIGA-CDG.

The *CNTN2* variant identified in the pedigree and the *in vivo* testing makes *CNTN2* a strong candidate genetic modifier of *PIGA* and may explain the incomplete penetrance of PIGA-CDG. However, it is difficult to know for sure if it is the actual cause for the incomplete penetrance observed in the family. There were many other promising candidates that were eliminated with our focused pipeline. It is also possible that multiple variants are playing a role in causing incomplete penetrance of PIGA-CDG. We genotyped other members of the family, including the great-uncle’s children, to determine if there were others with the *PIGA^S132C^* variant that were unaffected; however, only one female member genotyped carried the *PIGA^S132C^* variant with none of the male members carrying the *PIGA^S132C^* variant. We also assumed that most of the variants were loss-of-function and used RNAi as our functional tests. It is possible that some might require over-expression. Over-expression analyses are much larger experiments that require generating many new transgenic stocks and will be an important focus for future analyses. While we cannot definitively state that the incomplete penetrance of PIGA-CDG in this family is due to a variant in the *CNTN2* gene, we do show that loss of *CNTN2* function can improve *PIGA* loss of function phenotypes.

Incomplete expressivity and reduced penetrance remain large problems in the genetics of rare diseases. Both are likely more prevalent than currently appreciated. In human genetics, identifying pedigrees that show either phenomenon is difficult. This is especially challenging in rare diseases such as PIGA-CDG, where large non-consanguineous pedigrees are rare. However, identifying pedigrees with these phenomena is key to identifying modifier genes. By working closely with PIGA-CDG patients, family members, clinicians, and basic scientists, we were able to perform a pedigree analysis that identified multiple potential candidate genetic modifiers of *PIGA*. *In vivo* testing in fast, efficient models like *Drosophila* allowed for testing many candidates and quickly identifying *CNTN2* as a promising genetic modifier of PIGA-CDG. This study is one of the first to use pedigree information from patients along with an *in vivo* system to identify genetic modifiers of a rare disease. Further work between families, clinicians, and basic scientists can better enable us to study rare diseases and identify pathology and treatment options for these under-studied rare disorders.

## Supporting information

Data S1: cdg_gpi.bed

Data S2: In vivo Drosophila eye measurments

Video S1: PIGA neuronal KD climbing assay.

Video S2: CNTN2 neuronal KD climbing assay.

Video S3: PIGA, CNTN2 neuronal KD climbing assay.

Video S4: PIGA glial KD bang sensitivity assay.

Video S5: CNTN2 glial KD bang sensitivity assay.

Video S6: PIGA, CNTN2 glial KD bang sensitivity assay.

## Acknowledgments

We thank the family for their generosity and participation in this study. We also thank Drs. Hugo Bellen and Oguz Kanka for generating and sharing the *PIG-A* null allele. This work was funded by NIGMS R35GM124780 and a grant from the Primary Children’s Hospital Center for Personalized Medicine, Salt Lake City, UT to CYC. HJT was supported by a NIDDK training grant T32DK1109660 at the University of Utah. BSP and ARQ were supported by NHGRI grants R01HG012252 and R01HG010757. MD was supported by the Developmental Biology Training Program T32HD007491-26A1 at the University of Utah. JLB was supported by the Bray Chair.

## Supplemental video legends

Video S1: *PIGA* neuronal KD climbing assay.

Video S2: *CNTN2* neuronal KD climbing assay.

Video S3: *PIGA, CNTN2* neuronal KD climbing assay.

Video S4: *PIGA* glial KD bang sensitivity assay.

Video S5: *CNTN2* glial KD bang sensitivity assay.

Video S6: *PIGA, CNTN2* glial KD bang sensitivity assay.

## Data and code availability

Code for variant calling is found in the Methods section. Bed file is included as Data S1 supplemental file. VCF and alias files available upon request

