## Supplemental Figures and Table for "Identification of *CNTN2* as a genetic modifier of PIGA-CDG through pedigree analysis of a family with incomplete penetrance and functional testing in *Drosophila*"

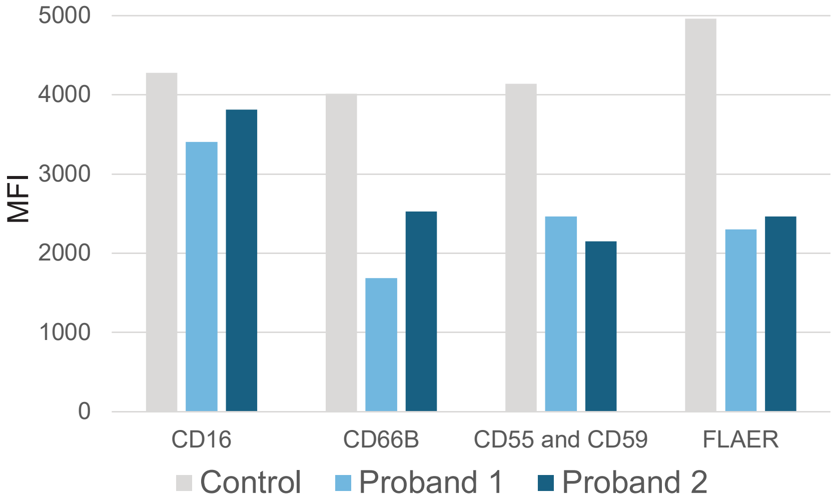


**Figure S1:** Measurement of GPI-anchored protein levels on the cell surface of control (grey) and proband (blue) cells. CD16, CD66B, CD55, and CD59 are specific GPI-anchored proteins. FLAER is a fluorescent protein binding all cell surface GPI-anchored proteins. MFI = mean fluorescence intensity.

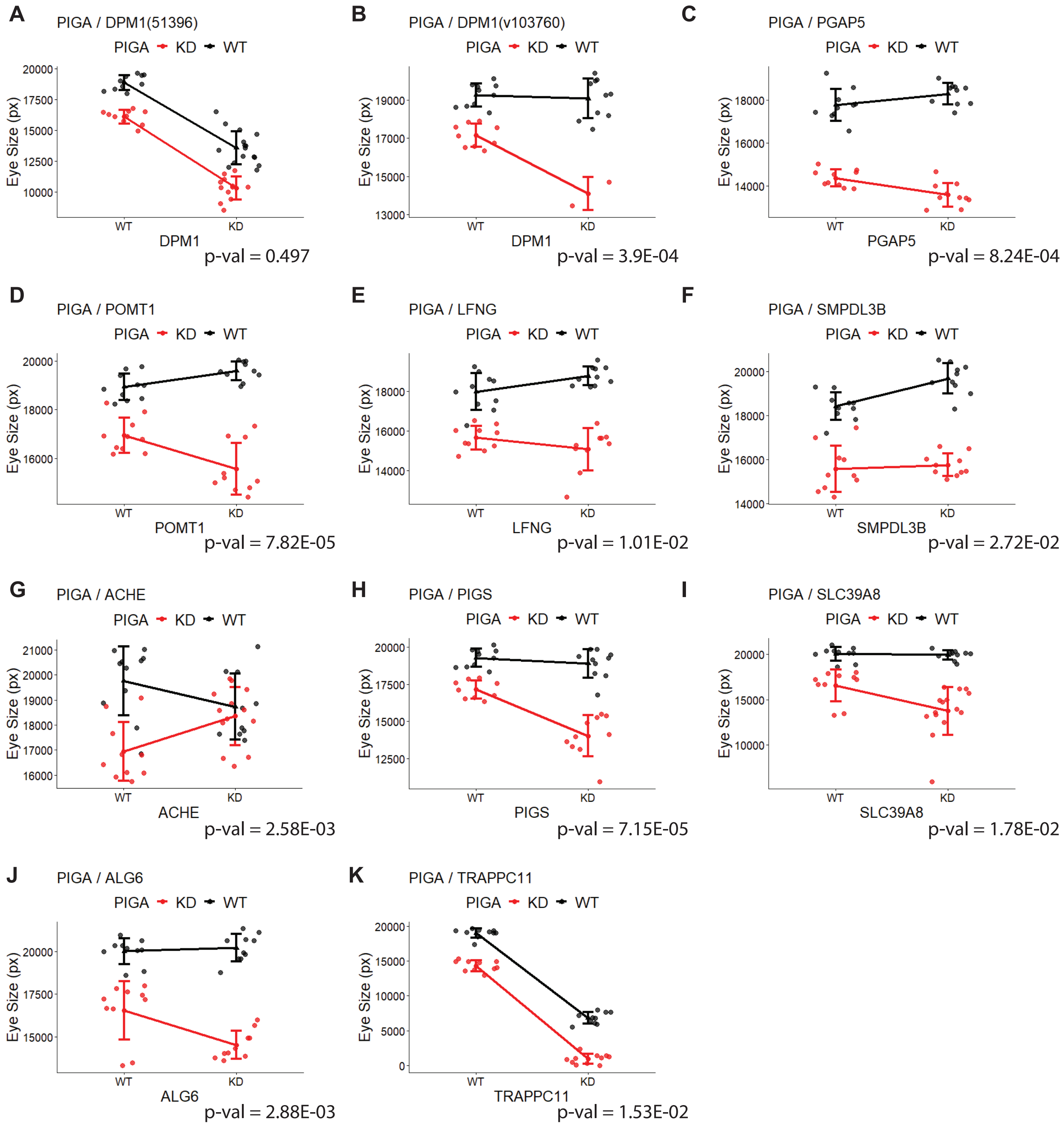


**Figure S2:** *In vivo* testing for genetic interactions in the eye. Quantification of eye sizes in pixels (px) is along the Y axis. X axis has expression level of candidate modifier protein, Wild Type (WT) or Knockdown (KD), and line color indicates expression level of PIGA (WT or KD). Interaction p-value calculated using a two-way ANOVA. Eye measurements found in supplemental file 2.

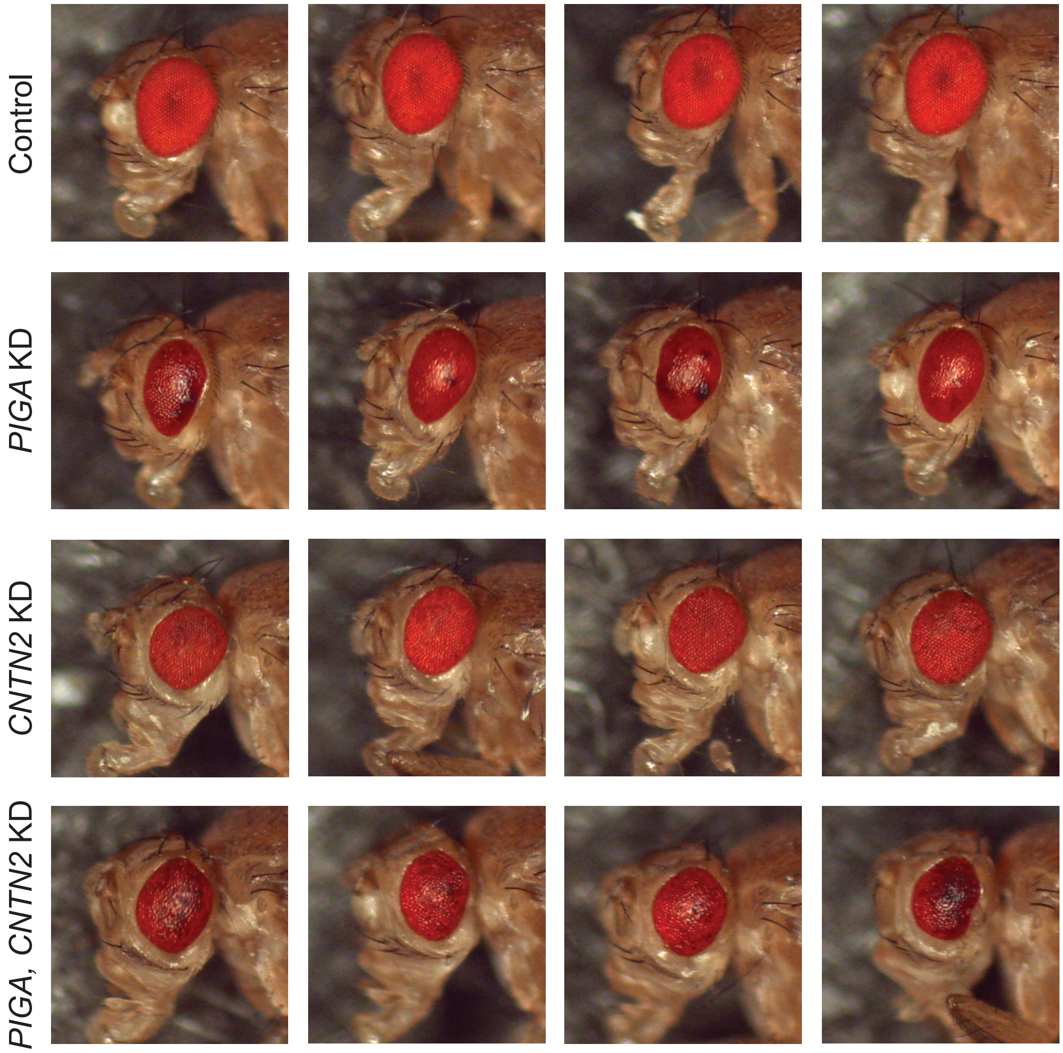


**Figure S3:** Representative images of *Drosophila* eyes showing Wild Type (WT) control, single Knockdown (KD), and double KD eyes for *PIGA* and *CNTN2*.

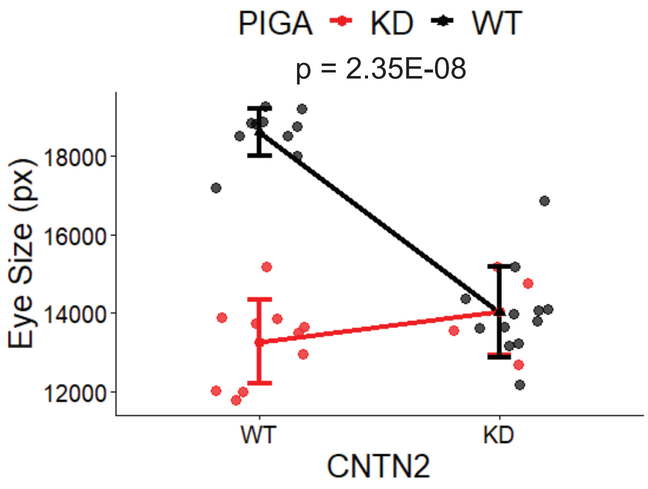


**Figure S4:** Genetic interaction between *PIGA* and *CNTN2* using a second RNAi. Quantification of eye sizes in pixels (px) is along the Y axis. X axis has expression level of candidate modifier protein. Wild Type (WT) or Knockdown (KD), and line color indicates expression level of *PIGA* (WT or KD). Parallel slopes indicate no interaction. Interaction p-value calculated using a two-way ANOVA.

**Table S1:** Number of variants and genes fitting each pipeline criterion

| Pipeline criterion | Protective | | Susceptibility | |
|---|---|---|---|---|
| | **Number of variants** | **Number of Genes** | **Number of variants** | **Number of genes** |
| Fit inheritance pattern | 701713 | 34770 | 787673 | 37260 |
| CDG gene/ GPI-anchored protein | 31 | 27 | 30 | 27 |
| Impact coding portion of genome | 19 | 18 | 24 | 23 |
| Predicted damaging by SIFT/PolyPhen | 7 | 7 | 13 | 13 |
